# Microglia initially seed and later reshape amyloid plaques in Alzheimer’s disease

**DOI:** 10.1101/2024.08.06.606783

**Authors:** Nóra Baligács, Giulia Albertini, Sarah C. Borrie, Lutgarde Serneels, Clare Pridans, Sriram Balusu, Bart De Strooper

## Abstract

We demonstrate the dual role of microglia in Alzheimer’s disease (AD), initially harmful, by seeding amyloid plaques, and protective in later stages by compacting amyloid plaques. Early microglial depletion using pharmacological or genetic blockage of CSF1R reduces plaque load and associated neuritic dystrophy, while human microglia transplantation restores plaque formation, confirming their seeding role. Transplanted *TREM2^R47H/R47H^*microglia exacerbate plaque pathology, highlighting microglia as key initiators of the amyloid pathology cascade.

## Introduction

Microglia respond to amyloid plaque accumulation in Alzheimer’s disease (AD), cluster around amyloid plaques, and change their transcriptional profiles adapting various cell states^1,2^. Many of the genetic risk genes associated with sporadic AD^3^ are expressed by microglia, even in their homeostatic state^2,4^. Microglia are generally believed to clear amyloid peptides and plaques from the brain^5^, but they are also implicated in chronic inflammation, neurotoxicity^6^, and the spread of amyloid and/or Tau pathology^7,8^. It remains unclear whether microglia act upstream in AD by increasing amyloid plaque formation (biochemical phase), during the disease’s cellular phase by modulating progression, or in the clinical phase by damaging synapses and neurons^9^, and whether the adaptive immunity affects their response in AD.

Many studies have reported either exacerbation, mitigation, or no change in amyloid plaque load following microglia depletion in AD mouse models (Supplementary Table 1). These conflicting results have been attributed to the different mouse models used, but we hypothesized that they might reflect different functions of microglia at different stages of amyloidosis. We therefore depleted microglia using PLX3397^10^ before amyloid plaque deposition, and after amyloid plaques have accrued. Additionally, we evaluated the role of peripheral adaptive immunity in the microglia-amyloid plaque interaction using *Rag2^-/-^* animals. We finally validated our observations using a genetic model of microglia depletion (*Csf1r^ΔFIRE/ΔFIRE^*)^11^. The findings indicate that microglia seed amyloid plaques in early disease and compact them at more advanced stages.

## Results

*App^NL-G-F^* mice were treated with PLX3397 in chow from 1 month (before plaque pathology) until 4 months of age. The amyloid burden and associated neuritic dystrophy were evaluated (Fig. 1). Microglia depletion was greater than 80%, as evaluated by IBA1^+^ staining (Extended Data Fig. 2a). At 4 months of age, both X34-positive amyloid fibrils and 82E1-immunoreactive Aβ deposits were reduced in the PLX3397-treated group. Microglia depletion reduced the area covered by amyloid plaques and the number of plaques but not their average size (Fig. 1b-d). ELISA measurements of soluble and insoluble Aβ brain fractions from the cortex confirmed this conclusion: soluble Aβ levels remained stable (apart from a minor decrease in Aβ38), while significant reductions of Aβ38, Aβ40, and Aβ42 were observed in the insoluble fraction (Fig. 1e). The reduction in amyloid plaques was accompanied by a reduction in LAMP1^+^ dystrophic neurites (Fig. 1f-g). Thus, microglia are seeding new plaques during the early amyloidosis phase in AD.

**Fig. 1:**
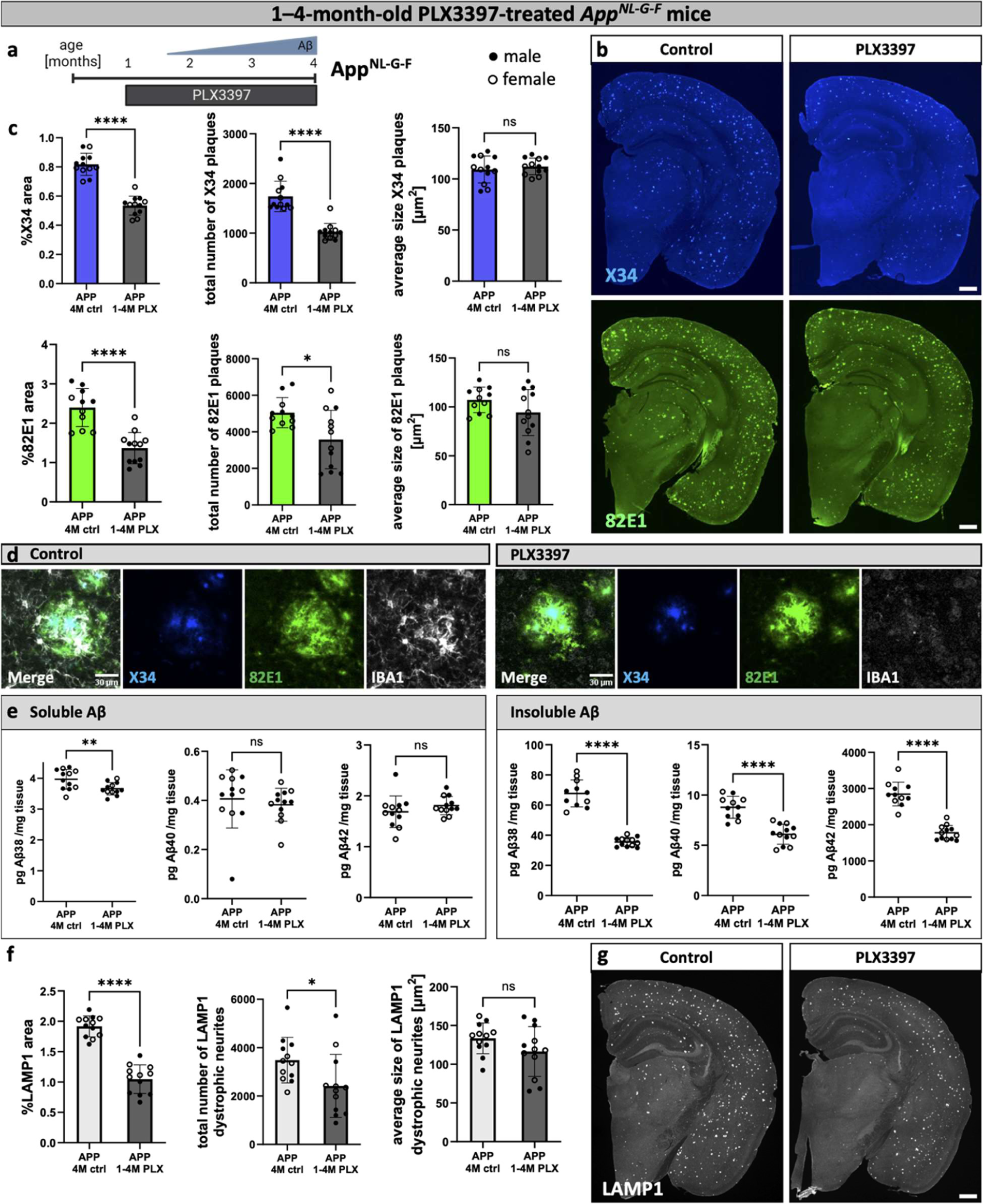
Microglia seed early amyloid plaques. (**a**) Treatment scheme. *App^NL-G-F^* mice (APP) were fed PLX3397 from 1 month until analysis at 4 months of age (1-4M PLX) or a control diet (4M ctrl). (**b**) Representative confocal images of amyloid plaques in the brain, stained with X34 (fibrillar plaques) or 82E1 (total Aβ). (**c**) Quantification of amyloid plaques in the whole brain (%X34 area: unpaired t-test, n(ctrl)=12, n(PLX)=12, p<0.0001; total number of X34 plaques: Mann-Whitney test, n(ctrl)=12, n(PLX)=12), p<0.0001; average size X34 plaques: unpaired t-test, n(ctrl)=12, n(PLX)=12, p=0.5098; %82E1 area: unpaired t-test, n(ctrl)=11, n(PLX)=12, p<0.0001; total number of 82E1 plaques: unpaired t-test with Welch’s correction, n(ctrl)=11, n(PLX)=12), p=0.012; average size 82E1 plaques: unpaired t-test, n(ctrl)=11, n(PLX)=12, p= 0.1225). (**d**) Higher magnification images of amyloid plaques and microglia in control and PLX3397-treated mice. (**e**) ELISA of amyloid levels in soluble and insoluble cortex extracts (sol. Aβ38: unpaired t-test, n(ctrl)=12, n(PLX)=12), p=0.0098; sol. Aβ40: Mann-Whitney test, n(ctrl)=12, n(PLX)=12), p=0.1725; sol. Aβ42: unpaired t-test, n(ctrl)=12, n(PLX)=12), p= 0.2571; insol. Aβ38: unpaired t-test with Welch’s correction, n(ctrl)=11, n(PLX)=12, p<0.0001; insol. Aβ40: unpaired t-test, n(ctrl)=11, n(PLX)=12, p<0.0001; insol. Aβ42: unpaired t-test, n(ctrl)=11, n(PLX)=12, p<0.0001). (**g**) Quantifications of dystrophic neurites around amyloid plaques (%LAMP1 area: unpaired t-test, n(ctrl)=12, n(PLX)=12), p<0.0001; total number of LAMP1 dystrophic neurites: unpaired t-test, n(ctrl)=12, n(PLX)=12), p=0.0317; average size of LAMP1 dystrophic neurites: unpaired t-test, n(ctrl)=12, n(PLX)=12), p=0.1321). (**f**) Representative images of LAMP1^+^ dystrophic neurites in the brain. White dots represent female mice and black dots represent male mice. Scale bars 500 μm (b, g), 30 μm (d, h). All data is presented as mean ± SD. *p ≤ 0.05; **p ≤ 0.01; ***p ≤ 0.001; ****p ≤ 0.0001.

Next, *App^NL-G-F^* mice were treated with PLX3397 after plaques were present, beginning at 3 months until 7 months of age (Extended Data Fig. 1a). In line with previous findings^12,13^, microglia depletion was less effective, however, IBA1 area was reduced by more than 60% (Extended Data Fig. 2b). 82E1-positive amyloid plaques increased significantly in area and size, but not in number (Extended Data Fig. 1b-d). The levels of soluble and insoluble Aβ in the brain were unchanged, except for a reduction in insoluble Aβ38 (Extended Data Fig. 1e). LAMP1-staining for dystrophic neurites was increased (Extended Data Fig. 1f-g). In conclusion, microglia do not alter the overall Aβ load in the brain but compact amyloid plaques and reduce dystrophic neurites in the late stage of amyloid plaque accumulation.

Sustained microglia depletion from before the onset of amyloid plaques (1 month) until 7 months of age (Extended Data Fig. 2d-h), resulted in a combination of the early seeding and late compacting effects of microglia on amyloid plaques. The total number of plaques was reduced, while the average plaque size was increased. Overall, this treatment regimen resulted in reduced amyloid area and insoluble Aβ levels in the brain (Extended Data Fig. 2e-g).

It has been suggested that the adaptive immune system plays a role in AD progression^14^. However, when we used a deletion of the *Rag2* gene to inhibit the generation of mature B- and T-lymphocytes^15^, we found a similar reduction of amyloid plaque size, number, and insoluble Aβ levels when microglia were depleted early (Extended Data Fig. 3a-c and 4a), and an increase in amyloid plaque area and size when microglia were depleted at more advanced stages of amyloidosis (3-7 months) (Extended Data Fig. 3d-f and 4b). Sustained microglia depletion (1-7 months) in the *App^NL-G-F^/Rag2^-/-^* mice (Extended Data Fig. 4c) also resulted in similar effects as observed in immunocompetent mice. Plaque numbers were reduced, while the average plaque size increased, resulting in a net decrease in plaque area and insoluble Aβ levels (Extended Data Fig.4c-g). Finally, the overall soluble and insoluble Aβ levels were similar in *App^NL-G-F^/Rag2^-/-^*mice compared to *App^NL-G-F^* mice across all treatments. This indicates that adaptive immunity does not alter the effect of microglia on amyloid plaque load or total amyloid levels in the *App^NL-G-F^* model.

We sought an independent way to confirm the effect of microglia on amyloid plaque pathology using mice that completely lack microglia because of a FIRE enhancer deletion (*Csf1r^ΔFIRE/ΔFIRE^*)^11^ (Extended Data Fig. 5a). We generated *App^NL-G-F^; Rag2^−/−^; IL2rg^−/−^; hCSF1^KI^; Csf1r^ΔFIRE/ΔFIRE^* mice to induce amyloid plaque pathology and allow the xenotransplantation of human microglia^2,16^ (hereafter referred to as FIRE mice). We xenotransplanted human-derived microglial progenitors^2,16^ into these mice at P4, and collected the brains 3 or 6 months after transplantation. Human transplanted microglia populate the entire mouse brain (Extended Data Fig. 5a). Additionally, staining with the disease-associated microglia (DAM)-marker CD9 is sparse and of low intensity in human microglia in 3-month-old FIRE mice (Extended Data Fig. 5b). By the age of 6 months, however CD9-staining is widespread and much more intense, where human CD9^+^ microglia also cluster around amyloid plaques (Extended Data Fig. 5c).

*App^NL-G-F^; Rag2^−/−^; IL2rg^−/−^; hCSF1^KI^* mice (control, with intact mouse microglia) show amyloid plaque deposition already at 6 weeks of age as demonstrated by 82E1 and X34 staining, which is not seen in FIRE mice (Fig. 2a). This result was confirmed by a 77% reduction in insoluble Aβ42 in the absence of microglia at the same age (Fig. 2b). By 3 months of age, FIRE mice develop some amyloid plaques, but to a much lower extent than control mice (Fig. 2c-e). Grafting of human microglia significantly restored plaque burden in the FIRE mice (Fig. 2c-e). Although the restoration is not complete, the result confirms the important contribution of microglia to amyloid plaque seeding at 3 months of age. By 6 months, the differences in amyloid load and dystrophic neurites between the control and FIRE mice groups are less pronounced but FIRE mice still have significantly reduced plaque load, insoluble Aβ levels (Extended Data Fig. 6a-c), and LAMP1^+^ dystrophic neurites (Extended Data Fig. 6d-e) compared to control mice. In contrast, levels of insoluble Aβ42 in xenotransplanted FIRE mice have reached similar levels as in control mice (Extended Data Fig. 6c).

**Fig. 2:**
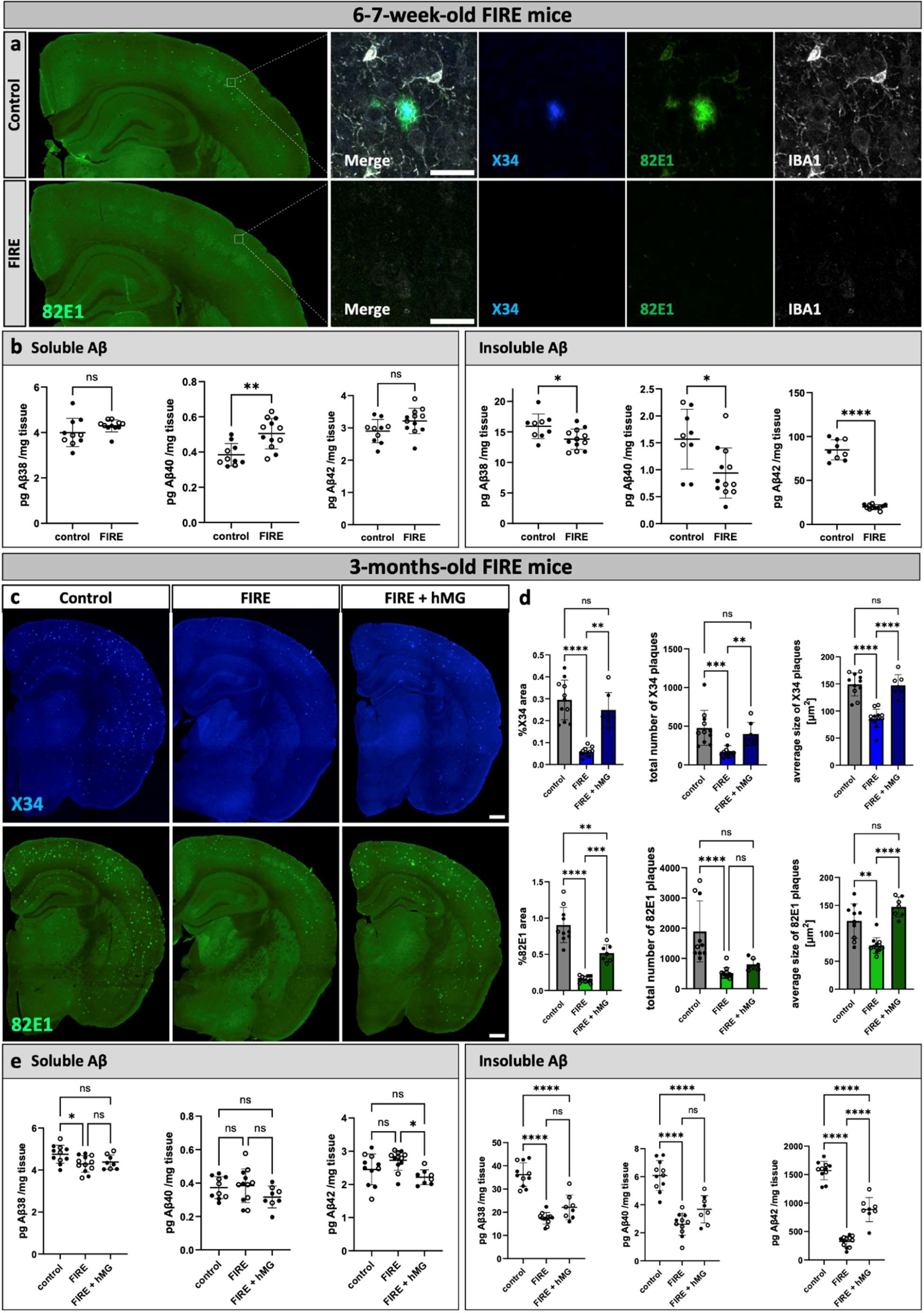
Genetic depletion of microglia delays amyloid plaque deposition and is reversed by xenotransplantation of human microglia. (**a**) Overview and higher magnification images of amyloid plaques in the brain and microglia in 6-7-week-old control mice (*App^NL-G-F^; Rag2^−/−^; IL2rg^−/−^; hCSF1^KI^,* with mouse microglia) and FIRE mice (*App^NL-G-F^; Rag2^−/−^; IL2rg^−/−^; hCSF1^KI^; Csf1r^ΔFIRE/ΔFIRE^*, no microglia). (**b**) Aβ ELISA of soluble and insoluble cortex extracts in 6-7-week-old control and FIRE mice (sol. Aβ38: unpaired t-test with Welch’s correction, n(ctrl)=10, n(FIRE)=11, p=0.1936; sol. Aβ40: unpaired t-test, n(ctrl)=10, n(FIRE)=12, p=0.0016; sol. Aβ42: unpaired t-test, n(ctrl)=10, n(FIRE)=12, p=0.0665; insol. Aβ38: unpaired t-test, n(ctrl)=9, n(FIRE)=12, p=0.017; insol. Aβ40: unpaired t-test, n(ctrl)=9, n(FIRE)=12, p=0.0107; insol. Aβ42: unpaired t-test with Welch’s correction, n(ctrl)=9, n(FIRE)=12, p<0.0001). (**c**) Representative images of amyloid plaques in the brains of 3-month-old control mice, FIRE mice, and FIRE mice xenografted with human microglia (FIRE + hMG), stained with X34 (fibrillar plaques) and 82E1 (total Aβ). (**d**) Quantifications of amyloid plaques in the whole brain (%X34 area: Welch’s ANOVA test, n(ctrl)=11, n(FIRE)=12, n(FIRE +hMG)=7, p<0.0001; total number of X34 plaques: Kruskal-Wallis test, n(ctrl)=11, n(FIRE)=12, n(FIRE +hMG)=7, p= 0.0001; average size X34 plaques: one-way ANOVA, n(ctrl)=11, n(FIRE)=12, n(FIRE +hMG)=7, p<0.0001; %82E1 area: Welch’s ANOVA test, n(ctrl)=10, n(FIRE)=12, n(FIRE +hMG)=7, p<0.0001; total number of 82E1 plaques: Kruskal-Wallis test, n(ctrl)=10, n(FIRE)=12, n(FIRE +hMG)=7, p<0.0001; average size 82E1 plaques: Welch’s ANOVA test, n(ctrl)=10, n(FIRE)=12, n(FIRE +hMG)=7, p<0.0001). (**e**) Aβ ELISA of soluble and insoluble cortex extracts (sol. Aβ38: one-way ANOVA, n(ctrl)=11, n(FIRE)=12, n(FIRE +hMG)=8, p=0.0237; sol. Aβ40: one-way ANOVA, n(ctrl)=10, n(FIRE)=12, n(FIRE +hMG)=8, p=0.1868; sol. Aβ42: one-way ANOVA, n(ctrl)=11, n(FIRE)=12, n(FIRE +hMG)=8, p=0.0125; insol. Aβ38: one-way ANOVA, n(ctrl)=11, n(FIRE)=12, n(FIRE +hMG)=8, p<0.0001; insol. Aβ40: one-way ANOVA, n(ctrl)=11, n(FIRE)=11, n(FIRE +hMG)=8, p<0.0001; insol. Aβ38: one-way ANOVA, n(ctrl)=11, n(FIRE)=12, n(FIRE +hMG)=8, p<0.0001). White dots represent female mice and black dots represent male mice. Scale bars 500 μm (a,c) and 30 μm (a). All data are presented as mean ± SD. *p ≤ 0.05; **p ≤ 0.01; ***p ≤ 0.001; ****p ≤ 0.0001.

Microglia are able to seed amyloid plaques before they adapt cell states associated with progressing AD (Extended Data Fig. 5b)^2,17^ and we therefore suggest that they act upstream in the classical amyloid cascade as the initial seeders of amyloid plaques. To test this further we grafted H9-derived WT human microglia and *TREM2^R47H/R47H^* microglia^18^ into the FIRE mice. TREM2 mutations impair microglia activation in response to amyloid pathology^2,19^. At 6-7 weeks of age, both WT and *TREM2^R47H/R47H^* xenotransplanted human microglia seeded amyloid plaques in FIRE mice (Extended Data Fig. 7), which were not present in non-grafted FIRE mice (Fig. 2a-c). The plaque load at that age was very low and not significantly different between WT and *TREM2^R47H/R47H^*microglia but the observation nevertheless reinforces the importance of microglia for the initial seeding of the amyloid plaques. Interestingly, at 3 months, insoluble Aβ42 levels were significantly increased in *TREM2^R47H/R47H^*microglia-grafted FIRE brains compared to WT microglia-grafted brains, suggesting increased seeding capacity of these microglia (Fig. 3c). Furthermore, the size of X34^+^ and 82E1^+^ plaques was increased as well, indicating in addition that the late protective function in plaque compaction is also affected in the *TREM2^R47H/R47H^* microglia (Fig. 3a,b). This was accompanied by an increase in LAMP1^+^ dystrophic neurite area (Fig. 3d-f). Thus *TREM2^R47H/R47H^*mutations exacerbate amyloid pathology both by increasing seeding of plaques in early amyloid deposition, and not being able to build the protective response once the plaques are formed.

**Fig. 3:**
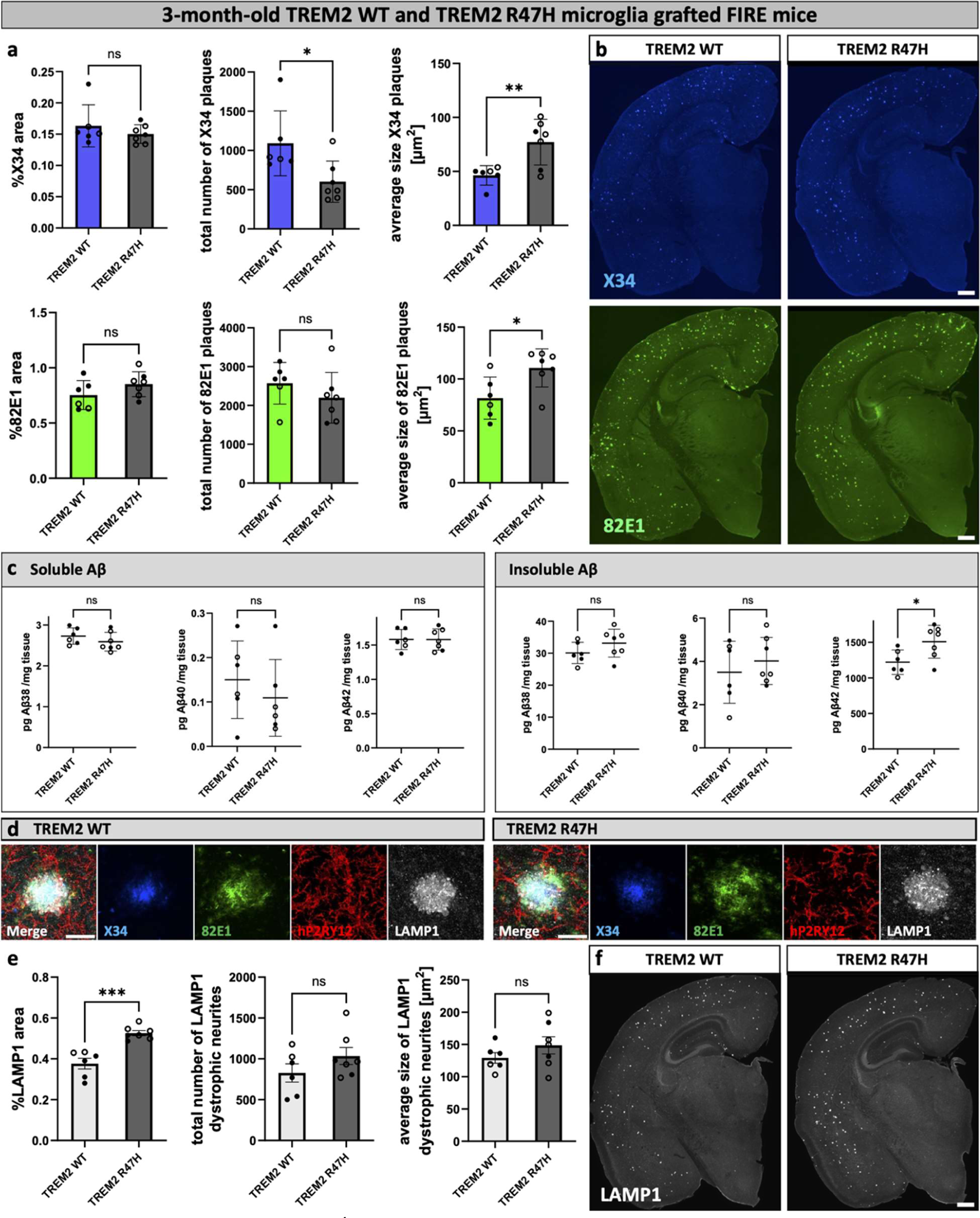
Xenotransplanted *TREM2^R47H/R47H^*microglia exacerbate amyloid and dystrophic neurite pathology in FIRE mice. FIRE mice (*App^NL-G-F^; Rag2^−/−^; IL2rg^−/−^; hCSF1^KI^; Csf1r^ΔFIRE/ΔFIRE^*) were xenotransplanted with human WT microglia (TREM2 WT) and human microglia harboring the *TREM2^R47H/R47H^*risk gene (TREM2 R47H) at P4 and analyzed at 3 months of age. (**a**) Image quantification of X34^+^ and 82E1^+^ amyloid plaques in the whole brain (%X34 area: Mann-Whitney test, n(TREM2 WT)=6, n(TREM2 R47H)=7, p=0.6282; total number of X34 plaques: Mann-Whitney test, n(TREM2 WT)=6, n(TREM2 R47H)=7, p=0.014; average size X34 plaques: unpaired t-test, n(TREM2 WT)=6, n(TREM2 R47H)=7, p=0.0072; %82E1 area: unpaired t-test, n(TREM2 WT)=6, n(TREM2 R47H)=7, p=0.1734; total number of 82E1 plaques: unpaired t-test, n(TREM2 WT)=6, n(TREM2 R47H)=7, p=0.2909; average size 82E1 plaques: unpaired t-test, n(TREM2 WT)=6, n(TREM2 R47H)=7, p=0.0201). (**b**) Representative images of amyloid plaques in the brain stained with X34 (fibrillar plaques) and 82E1 (total Aβ). (**c**) ELISA of amyloid levels in soluble and insoluble cortex extracts (sol. Aβ38: unpaired t-test, n(TREM2 WT)=6, n(TREM2 R47H)=7, p=0.2712; sol. Aβ40: unpaired t-test, n(TREM2 WT)=6, n(TREM2 R47H)=6, p=0.434; sol. Aβ42: unpaired t-test, n(TREM2 WT)=6, n(TREM2 R47H)=7, p=0.9832; insol. Aβ38: unpaired t-test, n(TREM2 WT)=6, n(TREM2 R47H)=7, p=0.1897; insol. Aβ40: unpaired t-test, n(TREM2 WT)=6, n(TREM2 R47H)=7, p= 0.4699; insol. Aβ38: unpaired t-test, n(TREM2 WT)=6, n(TREM2 R47H)=7, p= 0.0295). (**d**) Zoom in on plaques, dystrophic neurites, and human microglia staining in 3-month-old xenografted FIRE mice. (**e**) Quantification of LAMP1^+^ dystrophic neurites in the whole brain (%LAMP1 area: unpaired t-test, n(TREM2 WT)=6, n(TREM2 R47H)=7, p=0.0002; total number of LAMP1 dystrophic neurites: unpaired t-test, n(TREM2 WT)=6, n(TREM2 R47H)=7, p=0.2023; average size of LAMP1 dystrophic neurites: unpaired t-test, n(TREM2 WT)=6, n(TREM2 R47H)=7, p=0.2574). (**f**) Representative images of LAMP1^+^ dystrophic neurites in the whole brain. White dots represent female mice and black dots represent male mice. Scale bars 500 μm (b, f), 30 μm (d). All data is presented as mean ± SD. *p ≤ 0.05; **p ≤ 0.01; ***p ≤ 0.001; ****p ≤ 0.0001.

## Discussion

We examined the role of microglia in the amyloid cascade of AD, revealing a dual-phase response and putting homeostatic microglia upstream in the process. Early microglia depletion in the *App^NL-G-F^* model reduces plaque numbers without affecting their size, suggesting microglia’s involvement in seeding new plaques. After plaque formation, microglia depletion results in larger plaques but not an increased number, indicating microglia’s role in compacting plaques in later disease stages. In FIRE mice, the absence of microglia delayed the initiation of amyloid plaque pathology, while transplanting human microglia restored plaque pathology. The results demonstrate a critical role for the microglia in initiating amyloid plaques. Dystrophic neurite and amyloid staining closely correlate suggesting that the effect of microglia on dystrophic neurites is indirect by altering the amyloid plaque exposure of neurons (Extended Data Fig. 2c) The adaptive immune system does not influence microglia’s effect on amyloid plaque load.

Previous studies using various transgenic mouse models and microglia depletion methods have reported conflicting results. Our findings show that microglia’s impact on amyloid plaques varies with disease stages and cell state of the microglia, reconciling these discrepancies (Supplementary Table 1).

Despite the significant effects of microglia depletion on insoluble Aβ levels, soluble Aβ levels remained unchanged, indicating Aβ turnover is rather independent of microglia, which is unexpected. Therefore, plaque accumulation is not induced by increased soluble Aβ concentrations in our mice but is mediated by microglia, potentially through uptake, concentration^20^ and release^21,22^ of Aβ fibrils which seeds extracellular plaque formation.

We have previously shown that genetic risk of AD, assessed by the expression of associated risk genes, is spread over all different cell states of microglia, including homeostatic microglia^2^. Here, we provide observations that before microglia respond to plaques, they already carry the functional risk of initiating amyloid deposition putting them very early in the initiation of AD.

In conclusion, microglia are crucial in AD plaque deposition and pathology and our data emphasize the importance of considering disease stages and microglia activation states when targeting microglia for the treatment of AD.

## Extended Data Figures

**Extended Data Fig. 1:**
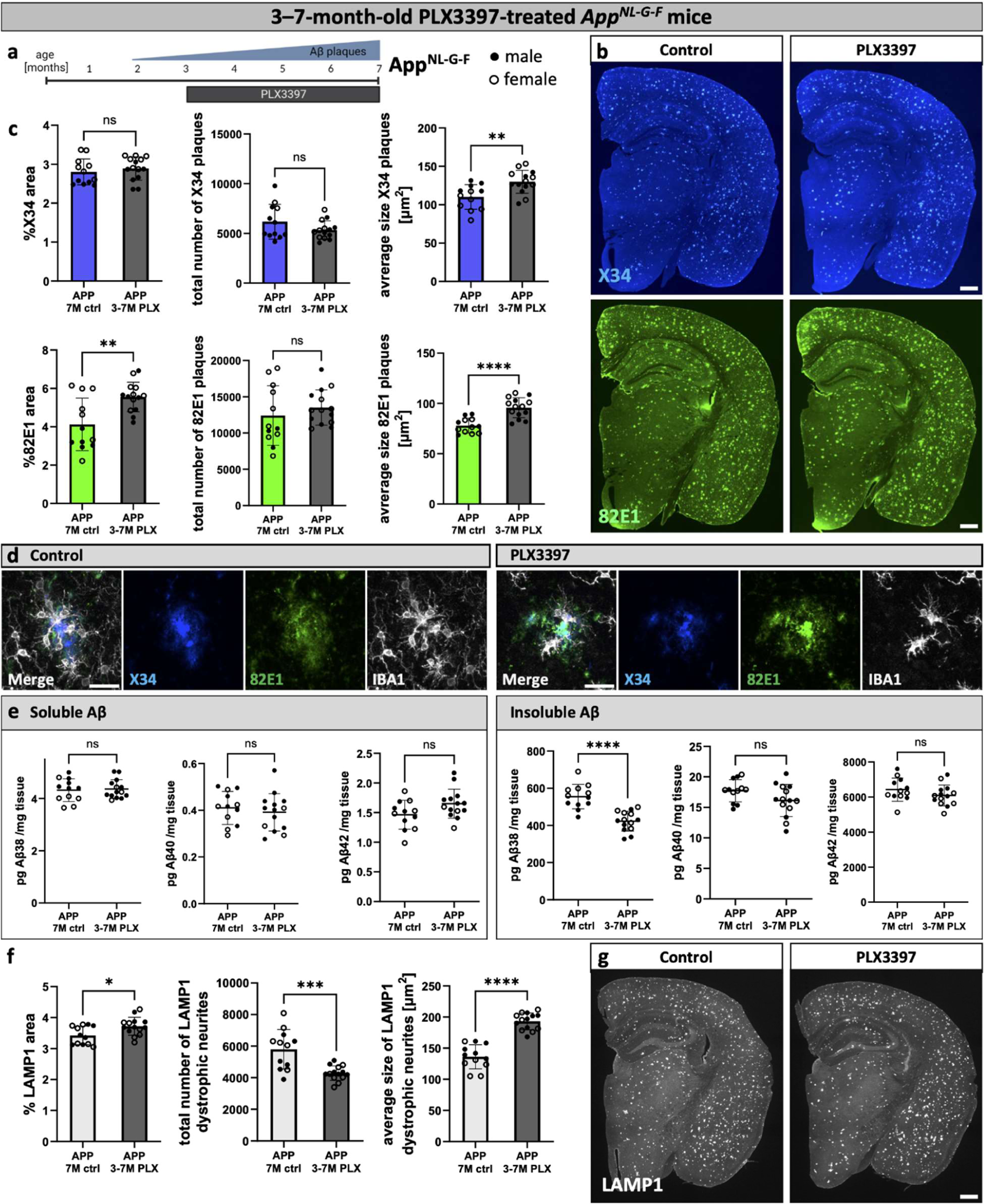
Microglia compact late amyloid plaques. (**a**) Treatment scheme. *App^NL-G-F^*mice (APP) were fed PLX3397 from 3 months until analysis at 7 months of age (3-7M PLX) or control diet (7M ctrl). (**b**) Amyloid plaques in the brain, stained with X34 (fibrillar plaques) and 82E1 (total Aβ). (**c**) Image quantifications of amyloid plaques in the whole brain (%X34 area: unpaired t-test, n(ctrl)=12, n(PLX)=14, p=0.4926; total number of X34 plaques: unpaired t-test with Welch’s correction, n(ctrl)=12, n(PLX)=14, p=0.1575; average size X34 plaques: unpaired t-test, n(ctrl)=12, n(PLX)=14, p=0.0034; %82E1 area: unpaired t-test, n(ctrl)=12, n(PLX)=14, p=0.0027; total number of 82E1 plaques: unpaired t-test, n(ctrl)=12, n(PLX)=14, p= 0.4042; average size 82E1 plaques: unpaired t-test, n(ctrl)=12, n(PLX)=14, p<0.0001). (**d**) Higher magnification images of amyloid plaques and microglia in control and PLX3397-treated mice. (**e**) ELISA of amyloid levels in soluble and insoluble cortex extracts (sol. Aβ38: unpaired t-test, n(ctrl)=12, n(PLX)=14, p=0.7964; sol. Aβ40: unpaired t-test, n(ctrl)=12, n(PLX)=14, p=0.5302; sol. Aβ42: unpaired t-test, n(ctrl)=12, n(PLX)=14, p=0.0638; insol. Aβ38: unpaired t-test, n(ctrl)=12, n(PLX)=14, p<0.0001; insol. Aβ40: unpaired t-test, n(ctrl)=12, n(PLX)=14, p=0.0887; insol. Aβ38: unpaired t-test, n(ctrl)=12, n(PLX)=14, p=0.1904). (**f**) Quantifications of dystrophic neurites around amyloid plaques (%LAMP1 area: unpaired t-test, n(ctrl)=12, n(PLX)=14, p=0.0172; total number of LAMP1 dystrophic neurites: unpaired t-test, n(ctrl)=12, n(PLX)=14, p=0.0003; average size of LAMP1 dystrophic neurites: unpaired t-test, n(ctrl)=12, n(PLX)=14, p<0.0001). (**g**) LAMP1^+^ dystrophic neurites in the brain. White dots represent female mice and black dots represent male mice. Scale bars 500 μm (b, g), 30 μm (d, h). All data is presented as mean ± SD. *p ≤ 0.05; **p ≤ 0.01; ***p ≤ 0.001; ****p ≤ 0.0001.

**Extended Data Fig. 2:**
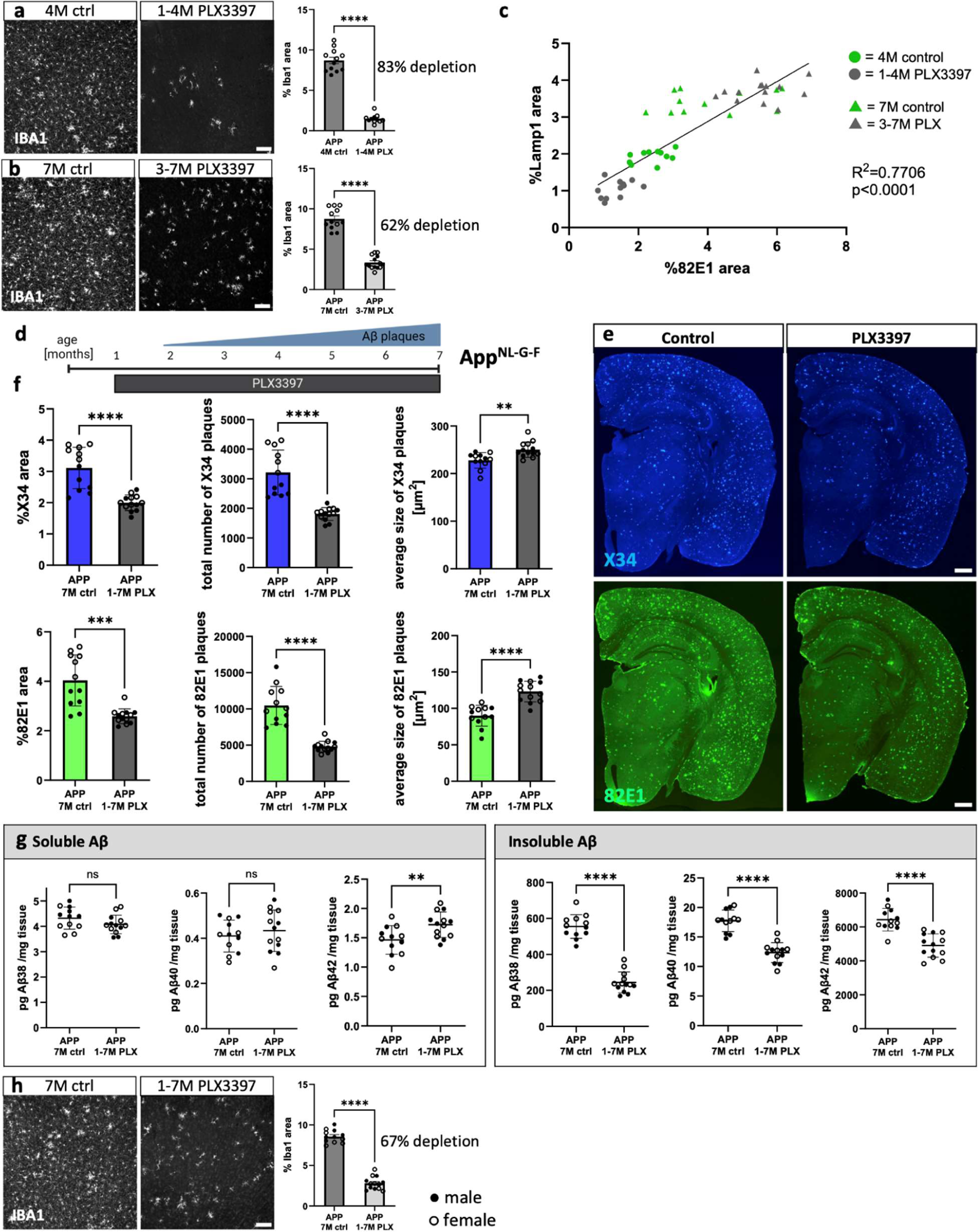
Additional information on PLX3397-treatment in *App^NL-G-F^* mice. (**a**) Representative images and microglia depletion efficiency of early PLX3397-treatment in *App^NL-G-F^* mice (APP) (unpaired t-test with Welch’s correction, n(ctrl)=12, n(PLX)=11, p<0.0001). (**b**) Representative images and quantification of microglia depletion efficiency of late PLX3397-treatment in *App^NL-G-F^* mice (unpaired t-test, n(ctrl)=12, n(PLX)=14, p<0.0001). (**c**) Linear regression of 82E1^+^ plaque staining and LAMP1^+^ dystrophic neurite staining across early and late microglia depletion cohorts (R^2^=0.7706, p<0.0001). (**d**) Treatment scheme for sustained microglia depletion. *App^NL-G-F^*mice (APP) were fed PLX3397 from 1 month until 7 months of age (1-7M PLX) or control diet (7M ctrl). (**e**) Representative images of amyloid plaques in the brain, stained with X34 (fibrillar plaques) and 82E1 (total Aβ). (**f**) Image quantifications of X34^+^ and 82E1^+^ plaques in the whole brain (%X34 area: unpaired t-test with Welch’s correction, n(ctrl)=12, n(PLX)=13, p<0.0001; total number of X34 plaques: unpaired t-test with Welch’s correction, n(ctrl)=12, n(PLX)=13, p<0.0001; average size X34 plaques: unpaired t-test, n(ctrl)=12, n(PLX)=13, p=0.0023; %82E1 area: unpaired t-test with Welch’s correction, n(ctrl)=12, n(PLX)=13, p=0.0005; total number of 82E1 plaques: unpaired t-test with Welch’s correction, n(ctrl)=12, n(PLX)=13, p<0.0001; average size 82E1 plaques: unpaired t-test, n(ctrl)=12, n(PLX)=13, p<0.0001). (**g**) ELISA of amyloid levels in soluble and insoluble cortex extracts (sol. Aβ38: unpaired t-test, n(ctrl)=12, n(PLX)=13, p=0.1392; sol. Aβ40: unpaired t-test, n(ctrl)=12, n(PLX)=13, p=0.4911; sol. Aβ42: unpaired t-test, n(ctrl)=12, n(PLX)=13, p=0.0098; insol. Aβ38: unpaired t-test, n(ctrl)=12, n(PLX)=13, p<0.0001; insol. Aβ40: unpaired t-test, n(ctrl)=12, n(PLX)=13, p<0.0001; insol. Aβ38: unpaired t-test, n(ctrl)=12, n(PLX)=13, p<0.0001). (**h**) Representative images and quantification of microglia depletion efficiency after sustained PLX3397 treatment (unpaired t-test, n(ctrl)=12, n(PLX)=13, p<0.0001). White dots represent female mice and black dots represent male mice. Scale bars 100 μm (a, b, h) and 500 μm (e). All data is presented as mean ± SD. *p ≤ 0.05; **p ≤ 0.01; ***p ≤ 0.001; ****p ≤ 0.0001.

**Extended Data Fig. 3:**
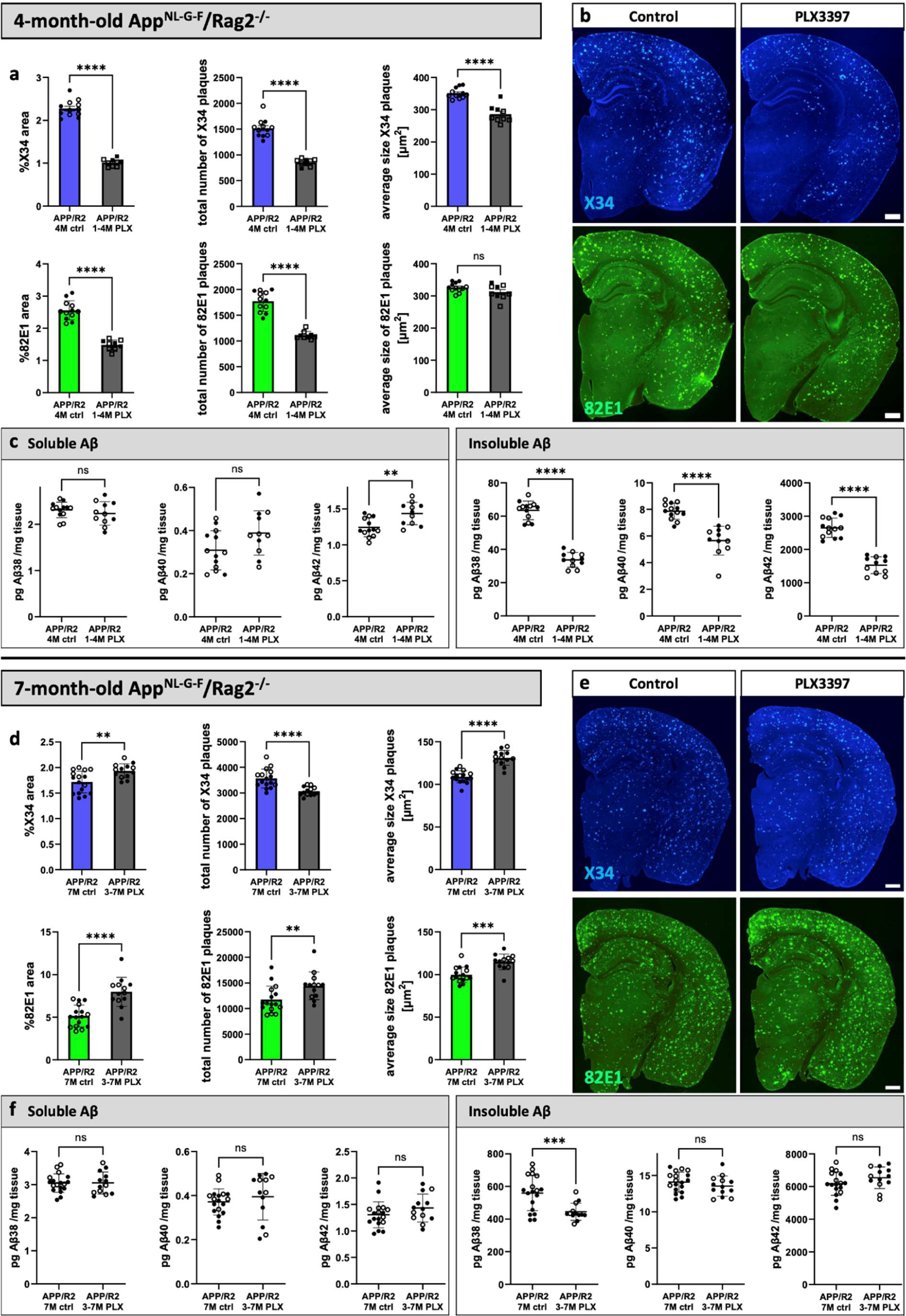
The adaptive immune system does not alter microglia-mediated modulation of amyloid plaques. (**a-c**) Immunodeficient *App^NL-G-F^/Rag2^-/-^*mice (APP/R2) were fed PLX3397 from 1 month until analysis at 4 months of age (1-4M PLX) or control diet (4M ctrl). (**a**) Quantifications of amyloid plaques in the whole brain of immunodeficient *App^NL-G-F^/Rag2^-/-^* mice (%X34 area: unpaired t-test with Welch’s correction, n(ctrl)=12, n(PLX)=9, p<0.0001; total number of X34 plaques: Mann-Whitney test, n(ctrl)=12, n(PLX)=9, p<0.0001; average size X34 plaques: unpaired t-test, n(ctrl)=12, n(PLX)=9, p<0.0001; %82E1 area: unpaired t-test, n(ctrl)=12, n(PLX)=9, p<0.0001; total number of 82E1 plaques: unpaired t-test with Welch’s correction, n(ctrl)=12, n(PLX)=9, p<0.0001; average size 82E1 plaques: unpaired t-test, n(ctrl)=12, n(PLX)=9,- p=0.0926). (**b**) Representative images of amyloid plaques in the brain, stained with X34 (fibrillar plaques) and 82E1 (total Aβ). (**c**) ELISA of amyloid levels in soluble and insoluble cortex extracts (sol. Aβ38: unpaired t-test, n(ctrl)=13, n(PLX)=11, p= 0.3824; sol. Aβ40: unpaired t-test, n(ctrl)=13, n(PLX)=11, p= 0.0558; sol. Aβ42: unpaired t-test, n(ctrl)=13, n(PLX)=11, p= 0.0029; insol. Aβ38: unpaired t-test, n(ctrl)=13, n(PLX)=11, p<0.0001; insol. Aβ40: Mann-Whitney test, n(ctrl)=13, n(PLX)=11, p<0.0001; insol. Aβ38: unpaired t-test, n(ctrl)=13, n(PLX)=11, p<0.0001). (**d-f**) Immunodeficient *App^NL-G-F^/Rag2^-/-^* mice (APP/R2) were treated with PLX3397 from 3 months until analysis at 7 months of age (3-7M PLX) or control diet (7M ctrl). (**d**) Image quantifications of amyloid plaques in the whole brain (%X34 area: unpaired t-test, n(ctrl)=16, n(PLX)=13, p=0.0036; total number of X34 plaques: unpaired t-test with Welch’s correction, n(ctrl)=16, n(PLX)=13, p<0.0001; average size X34 plaques: unpaired t-test, n(ctrl)=16, n(PLX)=13, p<0.0001; %82E1 area: unpaired t-test, n(ctrl)=16, n(PLX)=13, p<0.0001; total number of 82E1 plaques: Mann-Whitney test, n(ctrl)=16, n(PLX)=13, p=0.0056; average size 82E1 plaques: unpaired t-test, n(ctrl)=16, n(PLX)=13, p<0.0001). (**e**) Representative images of amyloid plaques in the brain of immunodeficient *App^NL-G-F^/Rag2^-/-^* mice, stained with X34 (fibrillar plaques) and 82E1 (total Aβ). (**f**) ELISA of amyloid levels in soluble and insoluble cortex extracts (sol. Aβ38: unpaired t-test, n(ctrl)=18, n(PLX)=13, p=0.9646; sol. Aβ40: unpaired t-test with Welch’s correction, n(ctrl)=18, n(PLX)=13, p=0.4748; sol. Aβ42: unpaired t-test, n(ctrl)=18, n(PLX)=13, p0.1756; insol. Aβ38: unpaired t-test with Welch’s correction, n(ctrl)=18, n(PLX)=13, p=0.0006; insol. Aβ40: unpaired t-test, n(ctrl)=18, n(PLX)=13, p=0.2682; insol. Aβ38: unpaired t-test, n(ctrl)=17, n(PLX)=13, p=0.1715). White dots represent female mice and black dots represent male mice. Scale bars 500 μm (a, d). All data is presented as mean ± SD. *p ≤ 0.05; **p ≤ 0.01; ***p ≤ 0.001; ****p ≤ 0.0001.

**Extended Data Fig. 4:**
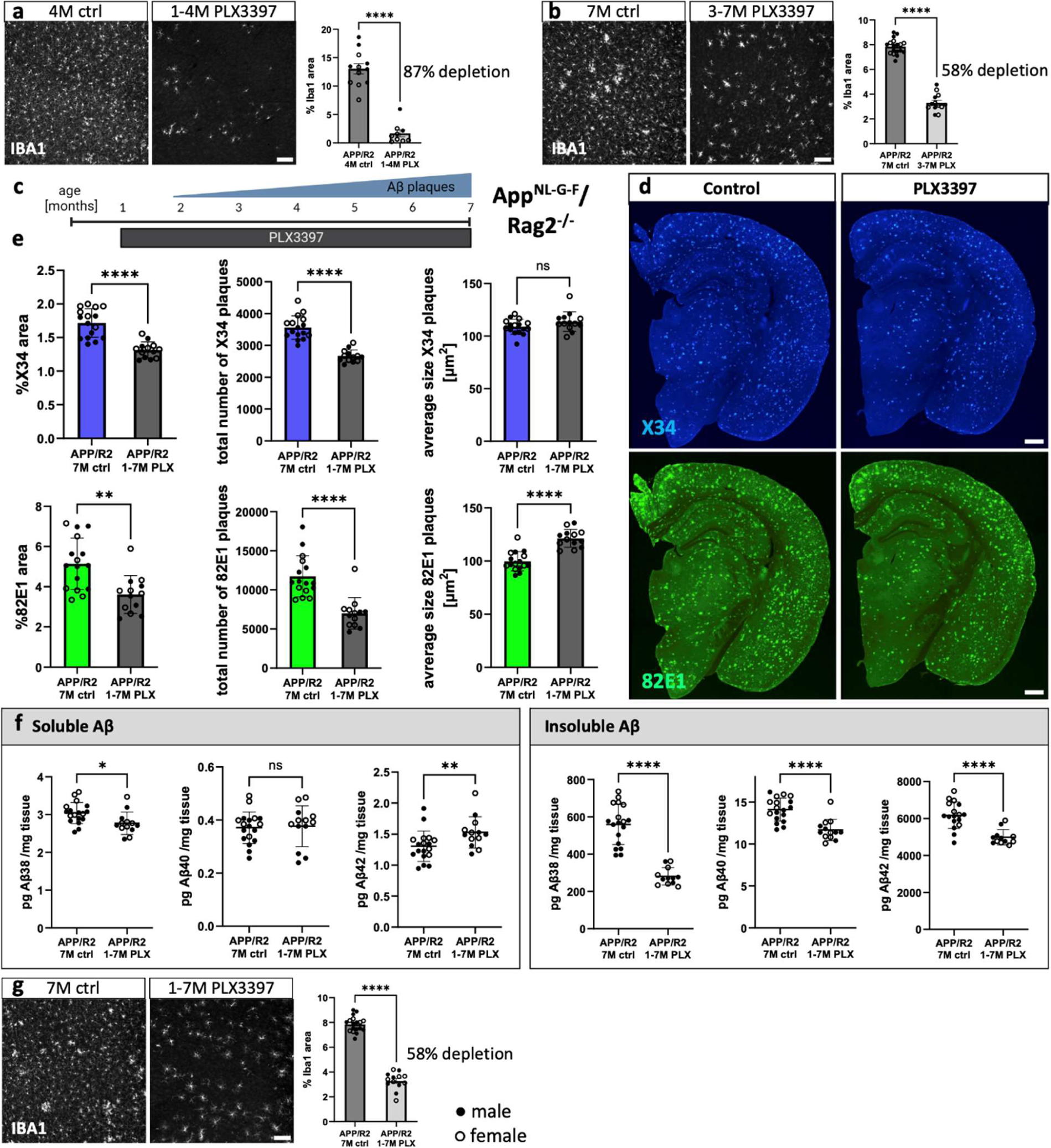
Additional information on PLX3397-treatment in *App^NL-G-F^/Rag2^-/-^* mice. **(a)** Microglia depletion efficiency of early PLX3397-treatment in *App^NL-G-F^/Rag2^-/-^*mice (APP/R2) (unpaired t-test, n(ctrl)=12, n(PLX)=9, p<0.0001)**. (b)** Representative images and quantification of microglia depletion efficiency of late PLX3397-treatment in *App^NL-G-F^/Rag2^-/-^*mice (APP/R2) (unpaired t-test, n(ctrl)=18, n(PLX)=13, p<0.0001). (**c**) Treatment scheme for sustained microglia depletion. *App^NL-G-F^/Rag2^-/-^*mice (APP/R2) were fed PLX3397 from 1 month until 7 months of age (1-7M PLX) or control diet (7M ctrl). (**d**) Representative images of amyloid plaques in the brain, stained with X34 (fibrillar plaques) and 82E1 (total Aβ). (**e**) Image quantifications of X34^+^ and 82E1^+^ plaques in the whole brain (%X34 area: unpaired t-test, n(ctrl)=16, n(PLX)=13, p<0.0001; total number of X34 plaques: unpaired t-test with Welch’s correction, n(ctrl)=16, n(PLX)=13, p<0.0001; average size X34 plaques: unpaired t-test, n(ctrl)=16, n(PLX)=13, p=0.1188; %82E1 area: unpaired t-test, n(ctrl)=16, n(PLX)=13, p=0.0013; total number of 82E1 plaques: Mann-Whitney test, n(ctrl)=16, n(PLX)=13, p<0.0001; average size 82E1 plaques: unpaired t-test, n(ctrl)=16, n(PLX)=13, p<0.0001). (**f**) ELISA of amyloid levels in soluble and insoluble cortex extracts (sol. Aβ38: unpaired t-test, n(ctrl)=18, n(PLX)=13, p=0.0146; sol. Aβ40: unpaired t-test, n(ctrl)=18, n(PLX)=13, p=0.8164; sol. Aβ42: Mann-Whitney test, n(ctrl)=18, n(PLX)=13, p=0.0073; insol. Aβ38: unpaired t-test with Welch’s correction, n(ctrl)=18, n(PLX)=12, p<0.0001; insol. Aβ40: unpaired t-test, n(ctrl)=18, n(PLX)=13, p<0.0001; insol. Aβ38: unpaired t-test, n(ctrl)=17, n(PLX)=12, p<0.0001). (**g**) Representative images and quantification of microglia depletion efficiency after sustained PLX3397 treatment (unpaired t-test, n(ctrl)=18, n(PLX)=13, p<0.0001). White dots represent female mice and black dots represent male mice. Scale bars 100 μm (a, b, g) and 500 μm (d). All data is presented as mean ± SD. *p ≤ 0.05; **p ≤ 0.01; ***p ≤ 0.001; ****p ≤ 0.0001.

**Extended Data Fig. 5:**
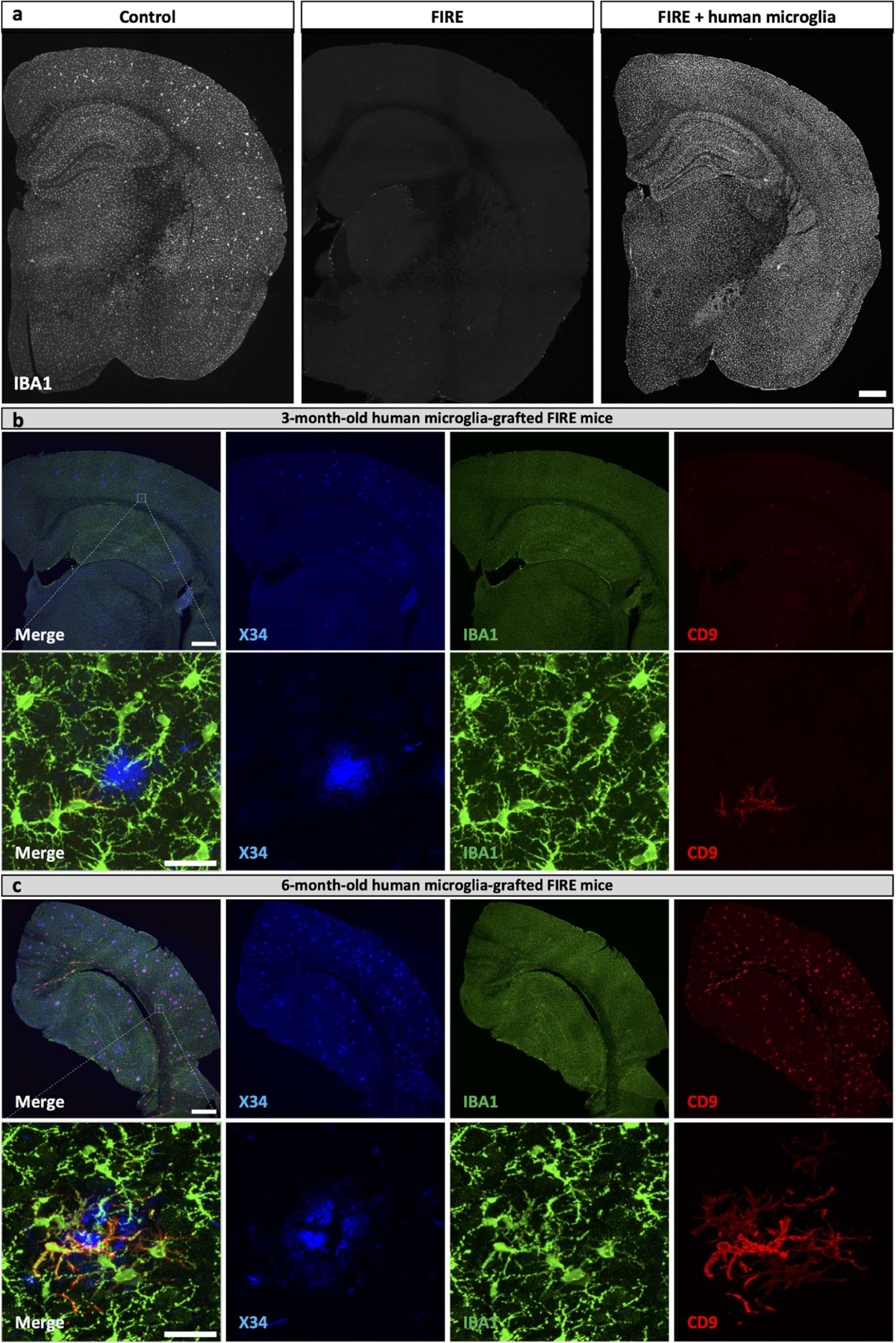
Human microglia grafted into FIRE mice populate the whole brain and upregulate the DAM marker CD9 around amyloid plaques. (**a**) Images of microglia in the brain of control mice (*App^NL-G-F^; Rag2^−/−^; IL2rg^−/−^; hCSF1^KI^,* with mouse microglia), FIRE mice (*App^NL-G-F^; Rag2^−/−^; IL2rg^−/−^; hCSF1^KI^; Csf1r^ΔFIRE/ΔFIRE^*, no microglia), and FIRE mice xenotransplanted with human microglia. (**b-c**) Overview and higher magnification images of human grafted microglia at 3 months (**b**) and 6 months (**c**) around amyloid plaques stained with IBA1 and the DAM marker CD9. Scale bars 500 μm (a,b,c) and 30 μm (b,c).

**Extended Data Fig. 6:**
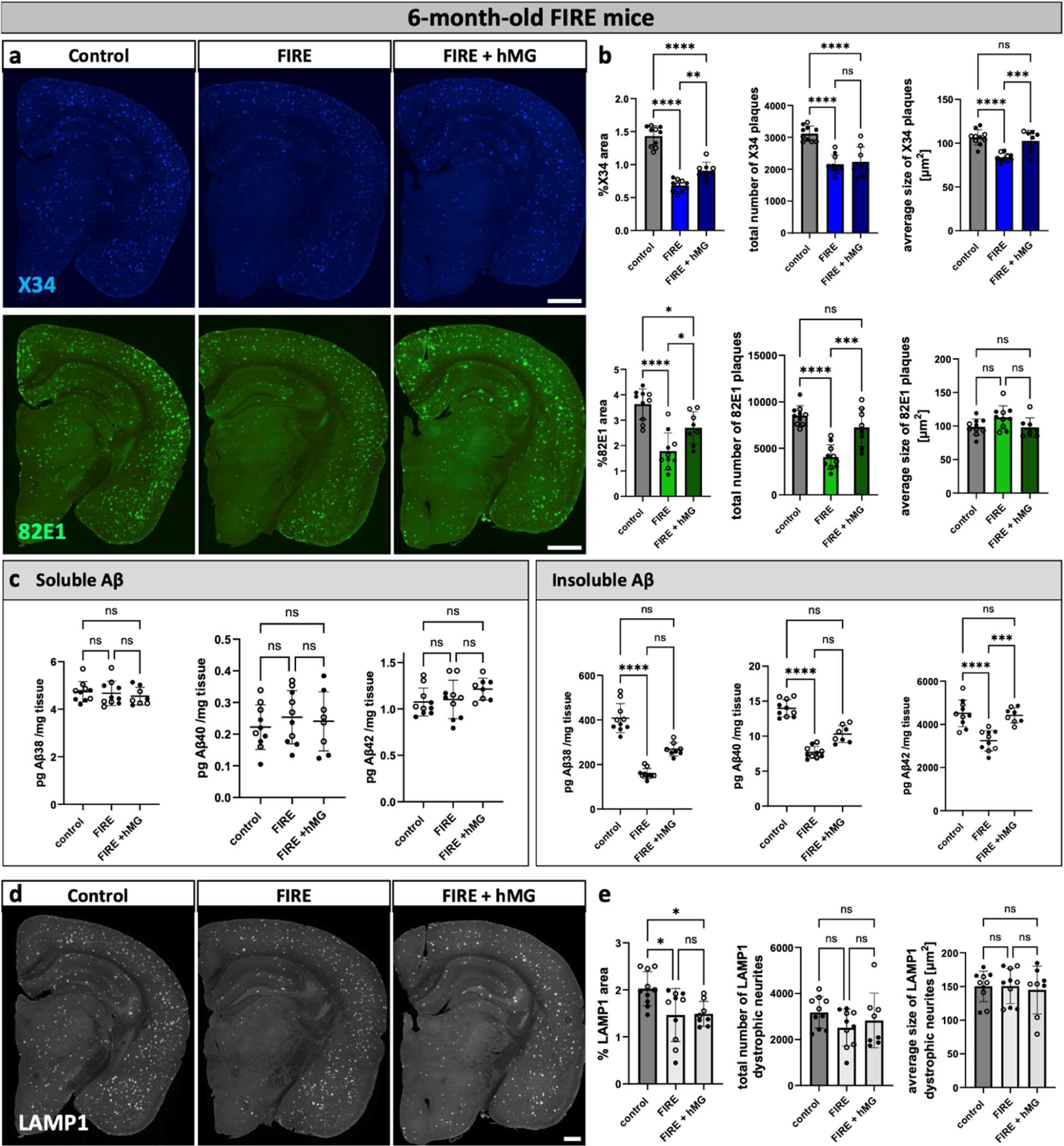
Reduced plaque burden in 6-month-old FIRE mice and rescue by human microglia transplantation. (**a**) Images of X34^+^ and 82E1^+^ amyloid plaques in the whole brain of control mice (*App^NL-G-F^; Rag2^−/−^; IL2rg^−/−^; hCSF1^KI^,* with mouse microglia), FIRE mice (*App^NL-G-F^; Rag2^−/−^; IL2rg^−/−^; hCSF1^KI^; Csf1r^ΔFIRE/ΔFIRE^*, no microglia), and xenografted FIRE mice (FIRE + hMG, human microglia). (**b**) Image quantifications of amyloid plaques in the whole brain (%X34 area: one-way ANOVA, n(ctrl)=10, n(FIRE)=10, n(FIRE +hMG)=8, p<0.0001; total number of X34 plaques: one-way ANOVA, n(ctrl)=10, n(FIRE)=10, n(FIRE+hMG)=8, p<0.0001; average size X34 plaques: one-way ANOVA, n(ctrl)=10, n(FIRE)=10, n(FIRE +hMG)=8, p<0.0001; %82E1 area: one-way ANOVA, n(ctrl)=10, n(FIRE)=10, n(FIRE +hMG)=8, p<0.0001; total number of 82E1 plaques: one-way ANOVA, n(ctrl)=10, n(FIRE)=10, n(FIRE +hMG)=8, p<0.0001; average size 82E1 plaques: one-way ANOVA, n(ctrl)=10, n(FIRE)=10, n(FIRE +hMG)=8, p=0.0594). (**c**) ELISA of soluble and insoluble Aβ levels in the cortex (sol. Aβ38: Kruskal-Wallis test, n(ctrl)=10, n(FIRE)=10, n(FIRE +hMG)=8, p=0.6434; sol. Aβ40: one-way ANOVA, n(ctrl)=10, n(FIRE)=10, n(FIRE +hMG)=8, p=0.6971; sol. Aβ42:m one-way ANOVA, n(ctrl)=10, n(FIRE)=10, n(FIRE +hMG)=8, p=0.2007; insol. Aβ38: Kruskal-Wallis test, n(ctrl)=10, n(FIRE)=10, n(FIRE +hMG)=8, p<0.0001; insol. Aβ40: Kruskal-Wallis test, n(ctrl)=10, n(FIRE)=10, n(FIRE +hMG)=8, p<0.0001; insol. Aβ38: one-way ANOVA, n(ctrl)=10, n(FIRE)=10, n(FIRE +hMG)=8, p<0.0001). (**d**) Images of LAMP1^+^ dystrophic neurites in the whole brain. (**e**) Image quantifications LAMP1^+^ dystrophic neurites in the whole brain (%LAMP1 area: one-way ANOVA, n(ctrl)=10, n(FIRE)=10, n(FIRE +hMG)=8, p=0.0118; total number of LAMP1 dystrophic neurites: one-way ANOVA, n(ctrl)=10, n(FIRE)=10, n(FIRE +hMG)=8, p=0.2678; average size of LAMP1 dystrophic neurites: Kruskal-Wallis test, n(ctrl)=10, n(FIRE)=10, n(FIRE +hMG)=8, p=0.8563). White dots represent female mice and black dots represent male mice. Scale bars 500 μm (a,d). All data is presented as mean ± SD. *p ≤ 0.05; **p ≤ 0.01; ***p ≤ 0.001; ****p ≤ 0.0001.

**Extended Data Fig. 7:**
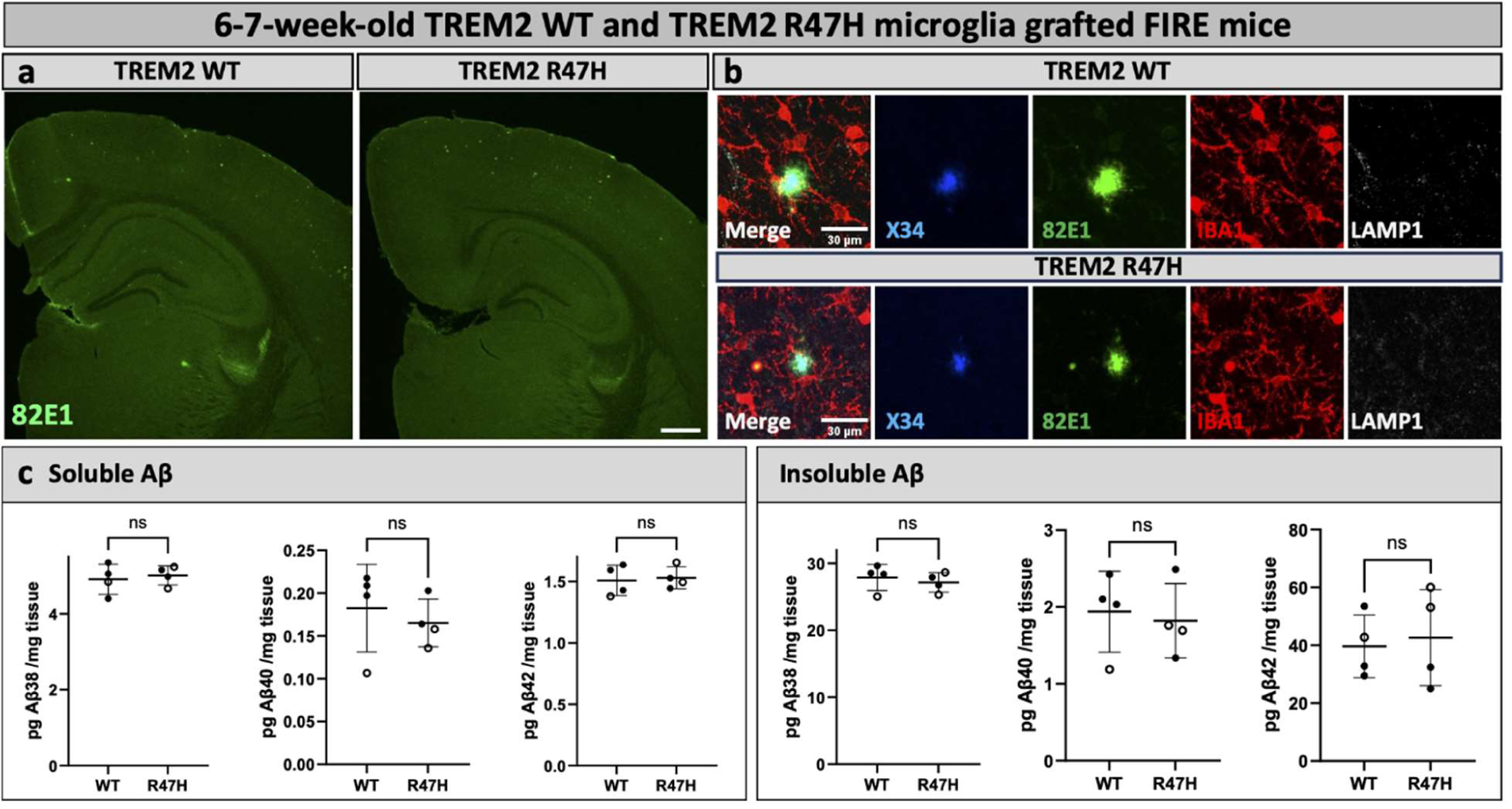
Amyloid pathology in 6-7-week-old FIRE mice transplanted with wildtype and *TREM2^R47H/R47H^*human microglia. FIRE mice (*App^NL-G-F^; Rag2^−/−^; IL2rg^−/−^; hCSF1^KI^; Csf1r^ΔFIRE/ΔFIRE^*) were xenotransplanted with human WT microglia (WT) and human microglia harboring the *Trem2^R47H/R47H^* (R47H) risk gene at P4 and analyzed at 6-7 weeks of age. (**a**) Overview images of amyloid plaques in the cortex of WT and R47H microglia grafted mice. (**b**) Higher magnification images on amyloid plaques, surrounded by human microglia. Early amyloid plaques do not yet show LAMP1^+^ dystrophic neurites. (**c**) ELISA of soluble and insoluble Aβ extracts of the cortex (sol. Aβ38: unpaired t-test, n(WT)=4, n(R47H)=13, p=0.6948; sol. Aβ40: unpaired t-test, n(WT)=4, n(R47H)=13, p=0.5737; sol. Aβ42: unpaired t-test, n(WT)=4, n(R47H)=13, p=0.7904; insol. Aβ38: unpaired t-test, n(WT)=4, n(R47H)=13, p=0.5686; insol. Aβ40: unpaired t-test, n(WT)=4, n(R47H)=13, p=0.7534; insol. Aβ38: unpaired t-test, n(WT)=4, n(R47H)=13, p=0.7714). White dots represent female mice and black dots represent male mice. Scale bars 500 μm (a) and 30µm (b). All data is presented as mean ± SD.

## Supplementary Table

**Supplementary Table 1:** Overview of microglia depletion studies in amyloid mice. This table summarizes the studies investigating the effect of microglia depletion on amyloid plaques. References, mouse model used, type of treatment and timing, microglia depletion efficiency, effect on amyloid plaques, and whether or not the results can be explained with our findings are indicated. Gray highlights indicate minor details that do not align with our observations.

| Study | Mouse model | Treatment | Depletion efficiency | Effect on plaque load | Explained |
| --- | --- | --- | --- | --- | --- |
| **APP/PS1** |  |  |  |  |  |
| Olmos-Alonso et al., 2016<sup>23</sup> | APP/PS1 | 6-9M<br>GW2580 | 30% | **Soluble Aβ ELISA:** <u>no change</u><br>**Histology:** <u>no change</u> in number of 6E10+ amyloid plaques in cortex | yes |
| Unger et al., 2018<sup>24</sup> | APP/PS1 | 12-13M<br>PLX5622 | 70-90% | **Histology:** <u>no change</u> in number or mean area of Thioflavin+ plaques in hippocampus and cortex | yes |
| Zhao et al., 2017<sup>5</sup> | *Cx3cr1<sup>CreER/+</sup>;R26<sup>DTR/+</sup>*<br>APP/PS1 | Diphtheria toxin at 15M, | 99%,<br>repopulation in 2<sup>nd</sup> week | **Intravital time-lapse imaging:** <u>no changes</u> in the number of Congo Red+ Aβ plaques, but an <u>increase in size</u> | yes |
| **5xFAD** |  |  |  |  |  |
| Spangenberg et al., 2019(Spangenberg et al., 2019) | 5xFAD | 1.5- 4M or 1.5-7M<br>PLX5622, repopulation | 97–100% | **Aβ ELISA:** no changes in soluble or insoluble Aβ or Aβ42<br>**Histology:** <u>reduced</u> Thio-S+ and 6E10+ plaque number and volume<br>Increased CAA in Cortex and thalamus | yes |
| Delizannis et al., 2021<sup>26</sup> | 5xFAD with AD-Tau injection | 1.5M-6M<br>PLX3397 | 92% | **Histology:** <u>reduced</u> NAB228+ Aβ signal in the cortex (layers 4-6), but not subiculum<br>Reduced plaque size in layers 4-6 of the cortex | yes |
| Sosna et al., 2018<sup>27</sup> | 5xFAD | 2-5M<br>PLX3397 | 99% | **Histology:** <u>reduction</u> of 6E10+ amyloid area and size | yes |
| Shabestari et al., 2022<sup>28</sup> | 5xFAD x FIRE | 0 – 5M or 0-6M | 100% | **Aβ ELISA:** <u>reduced</u> insoluble Aβ40 and 42, no change in soluble Aβ<br>**Histology:** <u>reduced</u> parenchymal plaques (AmyloGlo) in cortex, hippocampus, and Thalamus, increased CAA, | yes |
| Casali et al., 2020<sup>29</sup> | 5xFAD | 4-5M<br>PLX5622, repopulation | 50% | **Histology:** Slightly <u>reduced</u> 6E10 area in the thalamus, not in other brain regions<br><u>increased</u> diffuse-like plaques and fewer compact-like plaques | partially |
| Son et al., 2020<sup>30</sup> | 5xFAD | 9-10M<br>PLX3397 | 50% | **Histology:** <u>reduced</u> 6E10 Aβ deposition<br>**Western Blot:** decreased APP levels CTFβ, and Aβ in the cortex | no |
| Spangenberg et al., 2016(Spangenberg et al., 2016) | 5xFAD | 10-11M<br>PLX3397 | 90% | **Aβ ELISA:** <u>no change</u> in soluble or insoluble Aβ levels<br>**Histology:** <u>no change</u> in Thio-S+ and 6E10+ plaque numbers or size | yes |
| **App<sup>NL-G-F</sup>** |  |  |  |  |  |
| Benitez et al., 2021<sup>32</sup> | App<sup>NL-G-F</sup> | 1.5-3.5M<br>PLX5622 | nearly all microglia depleted | **Histology:** <u>reduction</u> in LCO+ plaque density of small plaques (hippocampus), | yes |
| Clayton et al., 2021<sup>33</sup> | App<sup>NL-G-F</sup> (with AAV-P301L-tau injection) | 4M-6M<br>PLX5622 | nearly all microglia depleted | **Histology:** <u>increased</u> Thio-S area, number of plaques, and average size of plaques<br>Increased 4G8 area and plaque size | yes |

## Methods

### Mice

*App^NL-G-F^* mice (*App^tm3.1Tcs^*)^34^ express amyloid precursor proteins (APP) at endogenous levels but contain the humanized Aβ sequence, as well as Swedish (NL; K670_M671delinsNL), Arctic (G; E693G), and Iberian (F; I716F) familial Alzheimer’s disease-causing mutations in C57BL/6 background. This strain was crossed with homozygous *Rag2* knockout mice (*Rag2^tm1.1Cgn^;* Jackson Laboratory, strain 008309) to generate *App^NL-G-F^/Rag2^-/-^*mice. To generate the FIRE mice, homozygous mouse oocytes from *Rag2^tm1.1Flv^; Csf1^tm1(CSF1)Flv^; Il2rg^tm1.1Flv^; App^tm3.1Tcs^* crosses^15^ in a mixed C57Bl6; BALB/c; 129S4 background were microinjected with reagents targeting the fms-intronic regulatory sequence (FIRE sequence) in the intron 2 of the mouse Csf1R gene as previously described by David Hume and Clare Pridans^11^. Ribonucleoproteins containing 0.3 μM purified Cas9HiFi protein 0.3 μM crRNA (5’GTCCCTCAGTGTGTGAGA3’ and 5’CAATGAGTCTGTACTGGAGC3’) and 0.3 μM trans-activating crRNA (Integrated DNA Technologies) were injected into the pronucleus of 120 embryos by the CBD Mouse Expertise Unit of KU Leuven. One female founder with the expected 428 bp deletion was selected and crossed with *a Rag2^tm1.1Flv^; Csf1^tm1(CSF1)Flv^; Il2rg^tm1.1Flv^;App^tm3.1Tcs^* male and the progeny were interbred to obtain *a Rag2^tm1.1Flv^; Csf1^tm1(CSF1)Flv^; Il2rg^tm1.1Flv^; App^tm3.1Tcs^*; *Csf1R^em1Bdes^*. For maintaining the colony *Rag2^-/-^; Csf1^CSF1/CSF1^; Il2rg^-/y^; App^NL-G-F/NL-G-F^; Csf1R^ΔFIRE/ΔFIRE^* males were crossed with *Rag2^-/-^; Csf1^CSF1/CSF1^; Il2rg^-/-^; App^NL-G-F/NL-G-F^; Csf1R^ΔFIRE/WT^* females as the 5 times homozygous females tend to take less care of their progeny. Mice were randomized, and both sexes were used for all experiments. Mice were housed in groups of two to five per cage with ad libitum access to food and water and a 14 h light/10 h dark cycle at 21°C. PLX3397 (Pexidartinib PLX3397, Asclepia MedChem Solutions), a CSF1R antagonist, was mixed in the mouse chow (600 mg/kg), which was replaced two times per week during the treatment period. All rodent experiments were approved by the Ethical Committee for Animal Experimentation (ECD) of KU Leuven and were executed in compliance with the ethical regulations for animal research.

### Immunofluorescence and Imaging

Mice were sacrificed with an overdose of sodium pentobarbital and immediately perfused with ice-cold 1 × DPBS (Gibco, Cat.#14190-144). After perfusion, one hemisphere was postfixed for 24 h at 4°C in 4% formaldehyde solution (PFA) for histology. The second hemisphere was dissected on ice. The olfactory bulb and cerebellum were removed, and the hippocampus, cortex, and midbrain were individually collected. The dissected brain regions were snap-frozen and stored at −80°C until use for biochemistry. For sectioning, the PFA fixed hemisphere was embedded in 4% Top Vision Low Melting Point Agarose (Thermo Scientific) and sectioned coronally into 35-40 µm sections using a vibrating microtome (Leica). Brain sections were collected under free-floating conditions and stored in cryoprotectant solution (40% PBS, 30% ethylene glycol, 30% glycerol) at −20 °C.

For staining, sections are washed in 1 × DPBS and permeabilized for 30 min at room temperature in PBS with 0.2% Triton. After permeabilization, sections were stained with X-34 staining solution (10 μM X-34 (Sigma-Aldrich), 20 mM NaOH (Sigma-Aldrich), and 40% ethanol) for 20 min at room temperature. X34 is a derivative of Congo red and is a fluorescent beta-sheet specific amyloid dye. Sections were washed three times with 40% ethanol for 2 min and twice with PBS + 0.2% Triton for 5 min. Sections were blocked with 5% normal donkey serum in PBS + 0.2% Triton X-100 for 1 h at room temperature. Primary antibody incubation was done overnight at 4 °C. The 82E1 antibody reacts with the N-terminal of amyloid Aβ peptides and, therefore, also stains diffuse Aβ-aggregates in the brain. The next day, sections were washed 3x with PBS and incubated with secondary antibodies for 2 h at RT. Sections were washed and finally mounted with Glycergel mounting medium (Agilent). Confocal images were obtained using a Nikon AX microscope with at ×4 (NA 0.2), ×20 (NA 0,75), x40 (NA 1.25), and x60 (NA 1.42). All images were acquired using similar acquisition parameters such as 16-bit, 1024×1024 quality, and images were processed in the FIJI/Image J software. All the images of Z-series stacks were then converted to Fiji/ImageJ maximum intensity projections.

For quantification of amyloid plaques (X34 and 82E1) and dystrophic neurites (LAMP1) in the whole brain hemisphere, large images with roughly 4-6 z-stacks were obtained with a 4x (NA 0.2) objective. Microscope settings were kept the same for all samples that were compared. Z-stacks were then converted to Fiji/ImageJ maximum intensity projections. The brain hemisphere was manually outlined with the polygon selection tool and added to the ROI manager. An intensity threshold for amyloid plaque quantification was selected manually and used for all the sections. In some sections, the threshold had to be adjusted, e.g., due to increased background. In those cases, a second threshold was determined which was applied for all the sections showing increased background. After thresholding, particles were analyzed and the relative area, number, and average size of particles were recorded. Two sections per brain were analyzed and the average values per brain were calculated. The same untreated 7-month-old mice were used as the control for the respective 3 to 7 months and the 1 to 7 months PLX3397 treatment. A few samples were excluded due to damaged sections or imaging artifacts. Excluded data is indicated in the source data.

For the quantification of microglia depletion, three 20x images of different cortical areas were acquired per mouse. Z-stacks were converted to maximum intensity projections and IBA1 staining was automatically thresholded (Otsu dark). Particles were analyzed to determine IBA1^+^ area.

| Antibody | Source | Identifier | dilution |
| --- | --- | --- | --- |
| Mouse anti-82E1 | IBL | 10323 | 1/200 |
| Rabbit anti-Iba1 | WAKO | 019-19741 | 1/200 |
| Guinea pig anti-Iba1 | Synaptic Systems | 234 308 | 1/500 |
| Rat anti-Lamp1 | Santa Cruz | sc-19992 | 1/500 |
| Rabbit anti-ubiquitin | Abcam | ab134953 | 1/250 |
| Mouse anti-human CD9 | BioLegend | 312102 | 1/500 |
| Rabbit anti-human P2RY12 | Atlas antibodies | HPA014518 | 1/500 |
| Donkey anti-mouse Alexa Fluor 488 | Invitrogen | A21202 | 1/500 |
| Donkey anti-rabbit Alexa Fluor 594 | Invitrogen | A21207 | 1/500 |
| Donkey anti-rat Alexa Fluor 647 | Abcam | ab150155 | 1/500 |
| Donkey anti-rabbit Alexa Fluor 488 | Invitrogen | A21206 | 1/500 |
| Donkey anti-mouse Alexa Fluor 594 | Invitrogen | A21203 | 1/500 |
| Donkey anti-guinea pig Cy5 | Jackson Immunolabs | 706-175-148 | 1/500 |

### Isolation of soluble and insoluble brain extracts and Aβ ELISA

Soluble and insoluble brain extracts were isolated from snap-frozen cortex samples. Samples were randomized and 8 samples across different cohorts were processed together. Snap-frozen cortices were thawed on ice and tissue weights were recorded. Ten volumes (w/v) P-TER buffer (Thermo Fisher, Cat.#78510) supplemented with cOmplete™ Protease Inhibitor Cocktail (Roche, Cat.#5056489001) was added and the tissue was homogenized in Fast prep tubes (MP Biomedicals) for 45 s at 6.5 m/s. Samples were centrifuged for 5 min at 5000 g to remove debris. Subsequently, the supernatant was centrifuged for 1h at 55,000 rpm at 4°C in an Optima Ultracentrifuge using a TLA110 rotor to pellet the insoluble brain fraction. The supernatant (=soluble fraction) was collected and stored at −80°C and the pellet was used for guanidine extraction.

Pellets were resuspended in 2 µl/mg tissue 6M GuHCl solution (6 M GuHCl/50 mM Tris-HCl, protease inhibitor cocktail, pH 7.6). Samples were sonicated with a micro-tip for 30 s at 10% amplitude, vortexed for 5 min, and incubated on a shaker for 1 h at 25°C and 450 rpm. Samples were ultracentrifuged for 20 min at 70,000 rpm and 4°C. The supernatant, containing guanidine-soluble Aβ fractions (insoluble Aβ) were transferred into a new tube, diluted 12 times with GuHCl diluent (20 mM phosphate, 0.4 M NaCl, 2 mM EDTA, 10% Block Ace, 0.2% BSA, 0.05% NaN3, 0.075% CHAPS, protease inhibitor cocktail, pH 7.0), and stored at −80°C until use.

Aβ38, Aβ40, and Aβ42 levels in the soluble and insoluble brain extracts were quantified by Meso Scale Discovery (MSD). Standard 96-well SECTOR plates (MSD, Cat.#L15XA-3) were coated with 1.5 µg/ml LTDA_38, LDA_40, or LTDA_Aβ42 capture antibody (homemade mouse monoclonal against Aβ_38_, Aβ_40_ or Aβ_42_ neoepitope respectively) in PBS, pH 7.4 at 4°C overnight. Plates were washed 5x with PBS-T (PBS + 0.05% Tween 20) and blocked with 0.1 % casein in PBS for 1.5 h at room temperature. Aβ standard curves were prepared with human Aβ1-38 (rPeptide Cat.#A-1078-1), Aβ1-40 (Cat.#A-1153-1), or Aβ1-42 (Cat.#A-1163-1). Samples were diluted according to previously determined concentrations. Diluted samples and standards were mixed 1:1 with LTDA_hAβN labeled with a sulfo-TAG detection antibody (homemade mouse monoclonal against the N-terminal sequence of human Aβ, in collaboration with Maarten Dewilde), in 0.1% casein in PBS, loaded on the blocked MSD plate, and incubated overnight at 4°C. Plates were washed 5x with PBS-T and 150 µl MSD GOLD Read Buffer A (Cat.#R92TG-2) was added to the wells. Plates were read with an MSD Sector Imager 2400A. Outliers were identified in GraphPad using ROUT (Q=1%) and excluded from analysis. Outliers were likely caused by technical issues, e.g., air bubbles or bad coating of individual wells. All data is available in the source data including excluded measures and reasons for exclusion.

### Differentiation of microglial progenitors and transplantation

H9-WT (WA09) human embryonic stem cells and H9 cells with *TREM2^R47H/R47H^*mutation were differentiated into microglia progenitors and transplanted into the brains of *App^NL-G-F^; Rag2^−/−^; IL2rg^−/−^; hCSF1^KI^; Csf1r^ΔFIRE/ΔFIRE^*mice following our published MIGRATE protocol^16^. *TREM2^R47H/R47H^* cells were acquired from the lab of Catherine M. Verfaillie at KU Leuven and have been generated from H9-WT (WA09) human embryonic stem cells as described by Claes et al^18^.

In brief, control and *TREM2^R47H/R47H^* H9-WA09 stem cells were plated and maintained on human Matrigel-coated six-well plates in E8 flex media until reaching ∼70–80% confluence. Once confluent, stem cell colonies were dissociated into single cells using Accutase (Sigma-Aldrich) and plated into U-bottom 96-well plates at a density of ∼10,000 per well in mTeSR1 medium with BMP4 (50 ng ml^−1^), VEGF (50 ng ml^−1^) and SCF (20 ng ml^−1^) for 4 days and allowed them to self-aggregate into embryoid bodies. On day 4, embryoid bodies were transferred into six-well plates (∼20 embryoid bodies per well) in X-VIVO (LO BE02-060F, Westburg) (+supplements) medium supplemented with SCF (50 ng ml^−1^), M-CSF (50 ng ml^−1^), IL-3 (50 ng ml^−1^), FLT3 (50 ng ml^−1^) and TPO (5 ng ml^−1^) for 7 days. A full medium change was performed on day 8. On day 11, the differentiation medium was replaced with X-VIVO (+supplements) with FLT3 (50 ng ml^−1^), M-CSF (50 ng ml^−1^) and GM-CSF (25 ng ml^−1^). On day 18, floating human microglial precursors were collected from the supernatant and engrafted into P4 mouse brains (0.5 million cells per pup) by bilateral injection with a Hamilton syringe as previously described^16^. Heterozygous *App^NL-G-F^; Rag2^−/−^; IL2rg^−/−^; hCSF1^KI^; Csf1r^+/ΔFIRE^* females and homozygous *App^NL-G-F^; Rag2^−/−^; IL2rg^−/−^; hCSF1^KI^; Csf1r^ΔFIRE/ΔFIRE^*males were used for breeding, or pups were fostered with CD1 mothers when breeding with *App^NL-G-F^; Rag2^−/−^; IL2rg^−/−^; hCSF1^KI^; Csf1r^ΔFIRE/ΔFIRE^* homozygous females to facilitate the survival. All experiments involving human stem cells have been approved by the UZ Leuven Ethical Committee and Biobank under study protocol S62888.

### Statistics and reproducibility

Statistical tests and data visualization were performed using GraphPad Prism10. Each data point represents one mouse and data is presented as mean± SD. Animals were randomly assigned to conditions to account for potential ordering effects. For all the ELISA data, statistical outliers (caused by technical errors) were identified using the ROUT test in Prism10 (Q=1%) and excluded from further analysis. 2-3 brain sections per mouse were analyzed and averaged for histological analysis. To avoid litter bias in the mouse experiments, experimental groups were composed of animals from different litters randomly distributed. Analysis was performed semi-automated or fully automated (when possible) using FIJI/Image J. Normality of residuals was checked with the Shapiro–Wilk tests and homoscedasticity was checked with the F-test or Brown-Forsythe test (for three groups). Comparisons between two groups following a normal distribution were analyzed using a two-tailed unpaired t-test, when homoscedasticity requirements were not met Welch’s correction was applied. Comparisons between two groups not following a normal distribution were analyzed with the Mann-Whitney test. When three groups were compared and data were normally distributed and homoscedastic, ordinary one-way ANOVA was used; when significant, it was followed by Tukey’s multiple comparisons test. When data were normally distributed but not homoscedastic, Welch’s ANOVA was used; when significant, it was followed by Dunnett’s multiple comparison. When data were not normally distributed, ranks were compared with Kruskal-Wallis followed by Dunn’s multiple comparisons test. Statistical significance was set at p<0.05. All data necessary for the conclusions of the study are available in the main text, figures, and Extended Data figures. Source data, including excluded data and reasons for exclusion, and detailed statistical analysis are provided with this paper.

## Acknowledgments

The authors thank Prof. Catherine Verfaillie for the *TREM2^R47H/R47H^* cells, Prof. Maarten Dewilde for the homemade antibodies used for Aβ ELISA’s. An Snellinx for help with the transplantation work. Amber Claes and Veronique Hendricks for breeding and taking care of the mice and the VIB Mouse Expertise Unit for generation of the Fire KO mice. This work was funded by a Medical Research Grant (MR/Y014847/1) awarded to B.D.S., the European Research Council (ERC) under the European Union’s Horizon 2020 Research and Innovation Programme (grant agreement no. ERC-834682 CELLPHASE_AD). The Flanders Institute for Biotechnology (VIB vzw), a Methusalem grant from KU Leuven and the Flemish Government, the Fonds voor Wetenschappelijk Onderzoek, KU Leuven, The Queen Elisabeth Medical Foundation for Neurosciences, the Opening the Future campaign of the Leuven Universitair Fonds, The Belgian Alzheimer Research Foundation (SAO-FRA) and the Alzheimer’s Association USA. B.D.S. holds the Bax-Vanluffelen Chair for Alzheimer’s Disease. N.B. is the recipient of a PhD fellowship from Fonds voor Wetenschappelijk Onderzoek (fellowship no. 11PSA24N). Schemes in Fig. 1a and Extended Data Figs. 1a, 2d and 4c were created with BioRender.com.

## Conflict of interests

B.D.S. has been a consultant for Eli Lilly, Biogen, Janssen Pharmaceutica, Eisai, AbbVie and other companies and is now consultant to Muna Therapeutics. B.D.S is a scientific founder of Augustine Therapeutics and a scientific founder and stockholder of Muna Therapeutics.

